# Integrin-Linked Kinase Controls Tissue Integrity in Embryonic Renal Stroma Via Expression of Fibronectin During a Critical Developmental Time Window

**DOI:** 10.64898/2026.11.20.768186

**Authors:** Xiangyue Hu, Norman D. Rosenblum

## Abstract

The stroma is a heterogenous framework that plays a critical role to mechanically and biologically support contiguous parenchymal elements in any organ. Studies on kidney development have historically focused largely on the epithelial and vascular tissue elements. Recent progress demonstrating stromal cell heterogeneity provided a basis to further investigate stromal functions at the crossroads of mechanical and chemical signaling in genetic mutants with stromal cell deficiency of Integrin-Linked Kinase, a Focal Adhesion (FA) adaptor, and a key regulator of the embryonic kidney. Our data demonstrate disruptions in stromal tissue integrity downstream of *Ilk* deletion, starting with loss of fibronectin and cellularity and stromal tissue tears, and eventually developing into fluid-filled tissue spaces with stromal cell death. We identified extracellular matrix (ECM) and FA programs as transcriptional and non-transcriptional targets of *Ilk* deletion and demonstrated that renal stroma normally gains these programs during development, which, when blocked by *Ilk* deficiency, leads to phenotypes that resemble a developmental arrest. Our data also identify E15.5-E16.5 as a critical developmental window for establishment of these programs.

## Introduction

The developing kidney arises from three major lineages: the nephrogenic lineage that develop into nephrons, the functional unit of filtration, the ureteric lineage that gives rise to the collecting system, and the stromal lineage that gives rise to a heterogenous supportive framework, including interstitial cells, vascular smooth muscle, pericytes, mesangial cells, cortical capsular cells, and fibroblasts (Rowan et al., 2017; Wilson & Little, 2021). Unlike the ureteric and nephrogenic lineages that are well-understood, investigations into the renal stroma have been obscured by its heterogeneity until recent advances made it possible to understand many biological processes at single-cell level (Magella et al., 2018; Lawlor et al., 2019; Naganuma et al., 2021; England et al., 2020; Combes et al., 2019). Spatial RNA sequencing of kidney sections throughout the course of kidney development (Jiang et al., 2025) then provided a spatial framework for understanding the transcriptome of stromal cells in relation to their location. These data provided the basis of understanding how stromal heterogeneity is established over the course of development, how stromal cells gain their biological functions in this heterogenous background, and more importantly, how these developmental processes can be regulated or disrupted by genetic regulators.

A key function of stromal cells is to provide structural support for developing multicellular organs. To this end, stromal cells produce the extracellular matrix (ECM), which maintains tissue integrity and holds signaling molecules required to regulated tissue development (Frantz et al., 2010). During this process, focal adhesions (FAs) are key components that translate these chemical and mechanical cues to regulate cellular processes (Mishra & Manavathi, 2021).

Integrin-Linked Kinase (ILK) is an adaptor protein at FAs. ILK acts to regulate cell-matrix interactions, cell adhesion, cell growth, apoptosis, and cell motility (Górska & Mazur, 2022). In the embryonic kidney, *Ilk* is known to be a key regulator in ureteric branching and podocyte and foot process architecture (Dai et al., 2006; El-Aouni et al., 2006; Smeeton et al., 2010; Smeeton et al., 2016). In the stromal lineage, while *Ilk* was found to play an essential role in *Pdgfrb*+ cells (Gong et al., 2018), which mainly consist of committed glomerular mesangial cells, vascular smooth muscle cells, and pericytes derived from stromal progenitors (Finer et al., 2026), the function of *Ilk* in stromal cell progenitors and their daughter interstitial cells is undefined.

Here, we investigate the role of *Ilk* in stromal development by generating mice with genetic deficiency of *Ilk* in embryonic kidney stromal cell progenitors and characterizing mutant tissue using single cell RNA sequencing. Our data demonstrate tissue disruption at the level of tissue, ECM, cells, and transcriptomes. By temporally characterizing these phenotypes, we reveal developmental changes of stromal cells underlying their biological function, which occurs at a critical developmental window. Our data demonstrate that loss of ILK blocks establishment of these developmental programs and leads to stromal tears in the outer medulla as kidney tissue expands over the course of development. Our data also suggest that fibronectin acts as a primary mediator downstream of ILK during this transition, which is regulated through non-transcriptional mechanisms.

## Results

### Kidneys with *Ilk*-deficient *Foxd1*+ stromal progenitors develop fluid-filled interstitial tissue spaces in the outer medulla

*Ilk* is globally expressed in the embryonic mouse kidney (Górska & Mazur, 2022). To understand the role of *Ilk* in development of the renal stroma, we introduced *Ilk* excision specific to *Foxd1*+ renal stromal progenitors using *Foxd1*-Cre (Humphreys et al., 2008) and *Ilk^fl/fl^* (Terpstra et al., 2003) mouse lines, generating *Foxd1^Cre/+^*;*Ilk^fl/-^* mutant mice. The number of *Foxd1^Cre/+^*;*Ilk^fl/-^* mice identified was consistent with the expected Mendelian ratio (1/4) during embryogenesis and at birth, but decreased postnatally, with a decreased number of mutants at P5 which remained viable as late as P12, indicating that stromal *Ilk* deletion impairs the viability of neonatal mice.

Mutant phenotypes were initially characterized at a post-natal stage. At post-natal day 0 (P0), *Foxd1^Cre/+^*;*Ilk^fl/-^*kidneys were characterized by medullary tissue spaces which can be shown, upon tissue dissection, to be fluid-filled (**Figure 1A**). Interestingly, although *Ilk* deficiency was introduced in the entire stromal lineage, the tissue spaces were only observed in the outer medullary region, and, as they expanded during kidney development, did not affect the cortical area (**Figure 1, A and C**).

**Figure 1:**
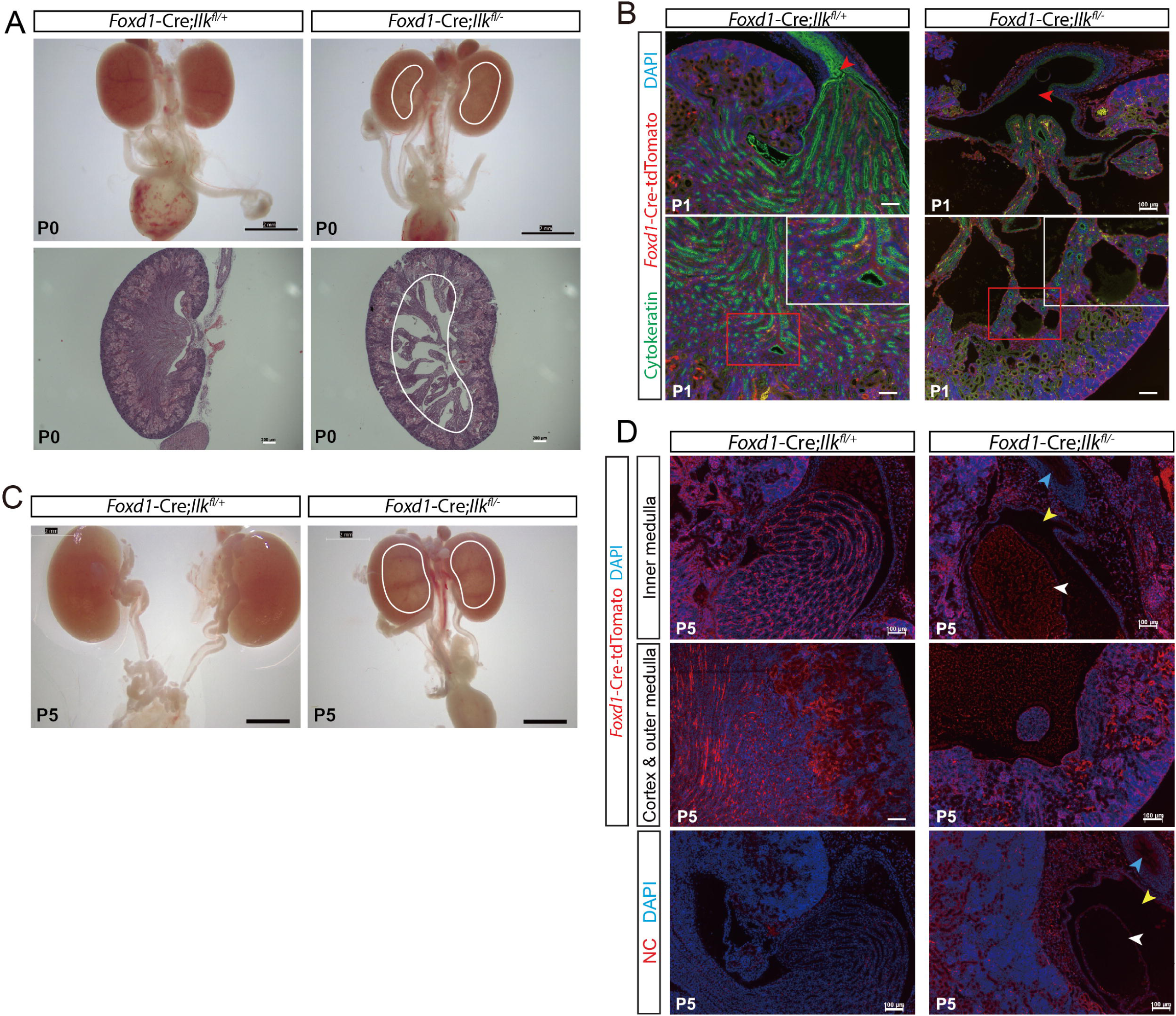
*Ilk* deletion in renal stromal progenitors results in fluid-filled tissue spaces in the medulla at post-natal stages. (A) Representative gross and histologic images of P0 kidneys. White lines highlight the outline of medullary tissue spaces in mutant kidneys. Scale bars: gross image: 2 mm; H&E staining: 200 μm. (B) Immunostaining of stromal tissue and epithelial tubules at P1. Red arrow heads highlight ureteropelvic junction (UPJ) tracts. No tdTomato-labeled, ectopic stromal cells were found to block the UPJ. Developing tissue spaces were lined by stromal cells as shown in the insets. Scale bars: 100 μm. (C) Gross images of P5 kidneys. White lines highlight the outline of tissue spaces in mutant kidneys. Cortical tissue remained grossly unaffected. Scale bars: 2 mm. (D) Immunostaining of stromal tissue at P5. NC: negative control sections processed without primary antibody. Arrow heads in the sections of *Ilk*-deficient kidney mark structures at the UPJ area. White: mutant renal papilla occupied by tissue space; yellow: renal pelvis; blue: ureter. Staining demonstrates presence of tdTomato+ cell debri that do not possess nuclei structures in the tissue space. Scale bars: 100 μm.

The observed fluid-filled tissue spaces are suggestive of hydronephrosis, a term referring to dilatation of the urinary collecting system (Cain et al., 2011). Previous studies have revealed mechanisms underlying the pathogenesis of hydronephrosis, including non-obstructive hydronephrosis caused by abnormal pacemaker function (Cain et al., 2011) in the ureter and obstructive hydronephrosis caused by ectopic stromal cells blocking the ureteropelvic (Sheybani-Deloui et al., 2018). To further understand the nature of fluid-filled interstitial spaces in *Ilk* mutant mice, immunofluorescence analysis was performed and showed normal ureteral development as indicated by normal expression of the smooth muscle and epithelial markers, SMA and Uroplakin, respectively **(Supplemental Figure S1A)**. To identify a possible blockage or narrowing in the urinary tract, intrapelvic dye injection was performed in postnatal kidneys (stages P1 and P5) and showed that dye injected into the mutant renal pelvis was able to enter the ureter and the bladder smoothly, indicating the absence of obstruction at the UPJ or in the ureter **(Supplemental Figure S1B)**. Together, these data indicate that fluid accumulation in medullary tissue is not due to hydronephrosis.

To further investigate the contribution of *Ilk*-deficient stromal cells in the formation of tissue spaces, *Rosa26-CAG-LSL-tdTomato* (referred to as *Rosa26^tdTm/tdTm^*) was introduced to label all descendant cells of the *Foxd1*+ stromal progenitors. Immunostaining at P1 confirmed that no ectopic tdTomato+ stromal cells are present at the UPJ in stromal-*Ilk*-deficient kidneys, further confirming the absence of UPJ obstruction (**Figure 1B**). Further, immunostaining revealed that the tissue spaces are lined by tdTomato-labeled interstitial cells, but not the typical Cytokeratin+ epithelial layer that characterize renal cysts (**Figure 1B**), suggesting that the tissue spaces arise in the interstitium rather than as tubular derivatives. Immunostaining at P5 also revealed the presence of tdTomato+ cell debris lacking nuclei staining in the tissue spaces (**Figure 1D**), suggesting that fluid that fills the tissue spaces contains dead cells that arise from the interstitium. Cell debris was limited to the tissue spaces, but was not observed in the renal pelvis and ureter, suggesting that tissue spaces do not represent dilated collecting ducts connected to the pelvis. Together, these findings suggest that malformation within *Ilk*-deficient interstitium causes tissue space formation in a cell-autonomous manner, which is related to interstitial cell death.

### Tissue space formation in stromal *Ilk*-deficient kidneys initiates at E16.5 with decreased medullary tissue expansion and stromal cellularity

To observe the formation of medullary tissue spaces, we next characterized *Ilk*-deficient kidneys histologically at progressive stages of embryonic development. While tissue spaces were present at E18.5 (**Figure 2A**), they could not be identified as early as E14.5 **(Supplemental Figure S2A)**, and E15.5 (**Figure 2A**) and were first observed at E16.5 when the interstitium could be seen to lose cellularity in the outer medulla (**Figure 2B**). Morphometric analysis of embryonic kidney tissue indicated that the medullary region comprises a mildly decreased portion of kidney area on mid-sagittal kidney sections in mutant kidneys. To better quantify this change, immunostaining was performed to detect LEF1, a marker specific for medullary interstitium (England et al., 2020), to visualize medullary interstitial cells and the boundary between medullary region and the renal cortex (**Figure 2D**). The area of medullary region was quantified in comparison to the entire area of the kidney. Data show that mutant kidneys exhibited a decrease in the portion of the medulla in the entire volume of the kidney starting at E16.5 (**Figure 2E**). As the overall geometry of the mutant kidneys appeared unchanged, this reduction suggests a shortened medulla and insufficient elongation of the epithelial tubules in the corticomedullary axis, suggestive of a non-cell-autonomous interaction in this region.

**Figure 2:**
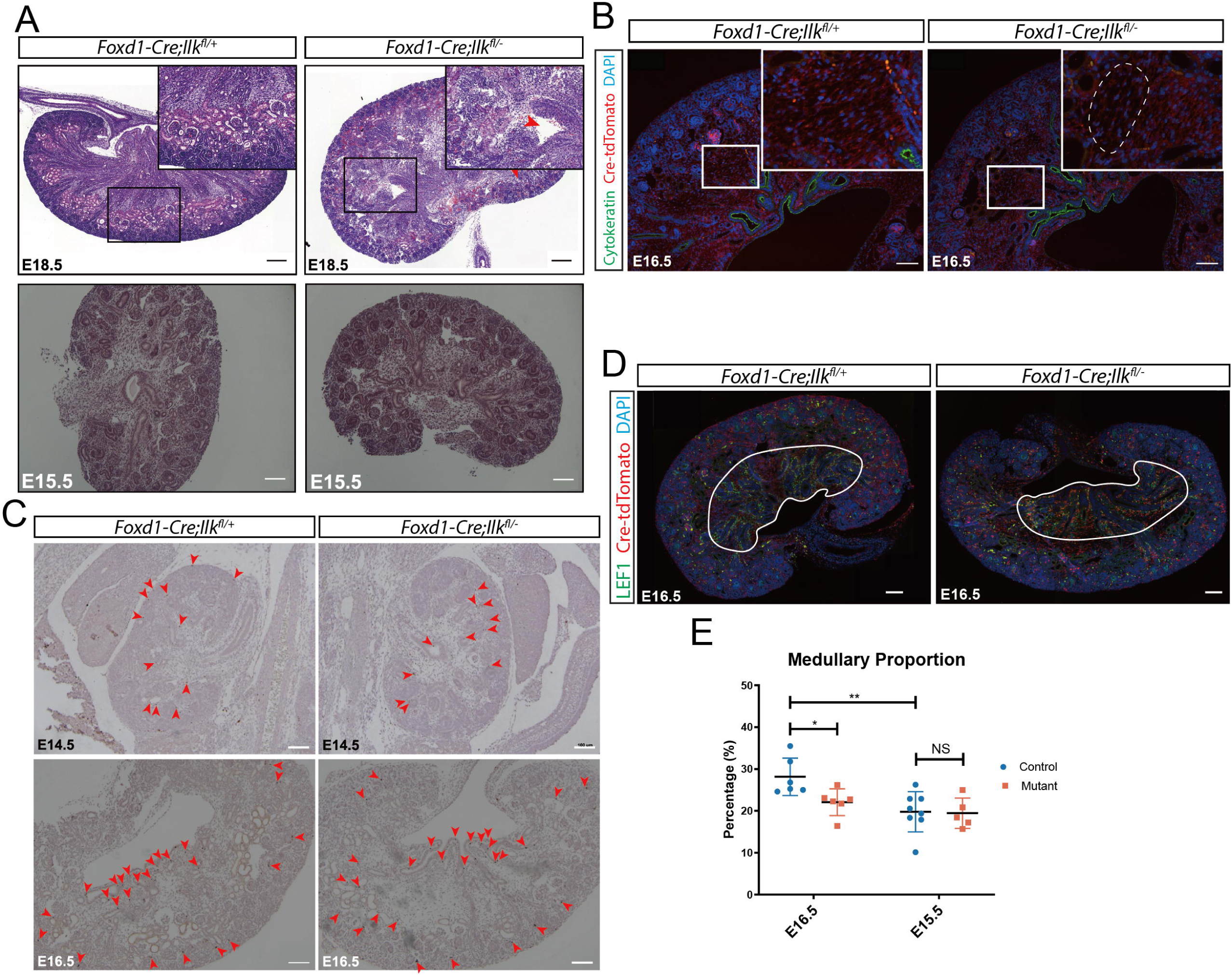
Fluid-filled tissue spaces arise from loss of cellularity in medullary stroma at E16.5 with no change in cell apoptosis. (A) Histologic observation on tissue space initiation in embryonic kidneys. Image shows presence of tissue spaces in the medulla of mutant kidneys at E18.5 as marked by the red arrowhead. Scale bars: E18.5: 200 μm; E15.5: 100 μm. (B) Early observations at E16.5 prior to formation of tissue spaces. White dashed line highlights sparsely populated outer medulla of mutant kidneys in comparison to control kidneys. Scale bars: 100 μm. (C) TUNEL assay performed on E16.5 and E14.5 kidneys. Red arrowheads highlight stained apoptotic cells. Data demonstrates no change in the number of apoptotic cells in the outer medulla at both timepoints. Scale bars: 100 μm. (D) E16.5 mutant kidneys demonstrated a decrease in medullary proportion of total kidney area at the mid-sagittal plane. White line highlights the boundary between the medulla and the cortex, as labeled by LEF1 staining. Scale bars: 100 μm. (E) Quantitation of the proportion of medullary area in total kidney area demonstrates a decrease in mutant kidneys that is only detectable at E16.5. Data are mean SD. * *p* < 0.05; ** *p* < 0.01 (two-way ANOVA test).

To test the hypothesis that tissue spaces arise via death of interstitial cells, a TUNEL assay was performed at E16.5, when the phenotypic abnormality in the medullary interstitium is first observed and before the spaces have developed, and at E14.5, earlier than any notable phenotypic abnormalities. At both stages, TUNEL assay showed no significant difference between *Ilk*-deficient kidneys and control kidneys in the pattern of cell apoptosis (**Figure 2C**). Apoptotic cells were located in the papillary and cortical region in both control and *Ilk*-deficient kidneys, while tissue space was observed to arise in the medullary region. Moreover, in both control and mutant kidneys, apoptotic cells were mostly concentrated in the epithelialized tubular populations and are rarely localized in the interstitium. We also tested cell proliferation by immunostaining for phospho-Histone H3, a label for proliferative cells. Our data demonstrated no significant difference in cell proliferation **(Supplemental Figure S2B)**. Like apoptotic cells, the proliferative cells also concentrated in papillary and cortical regions, not the phenotypic outer medulla. Together, these data suggest that interstitial cell death and decreased cell proliferation are not the cause of medullary tissue space formation.

### Single-cell RNA sequencing at E16.5 reveals matrisome and adhesion programs disrupted by stromal *Ilk* deficiency

Having identified E16.5 as the key timepoint for the onset of the phenotype, we further investigated the early transcriptional anomalies underlying the interstitial phenotype at this stage. Considering the heterogeneity of stromal tissue, single-cell RNA sequencing was performed to dissect the anomalies at a single-cell level. We identified 17 cell clusters from combined control and mutant cells, comprising ureteric, nephrogenic, stromal, and endothelial lineages (**Figure 3A**). Clusters were manually annotated by expression of population-specific markers of E16.5 kidneys (**Figure 3A, Supplemental Figure S3A)**. UMAP projection of cells showed a depletion of naïve nephron progenitors marked by *Cited1* expression (cluster 6) and an enrichment in collecting duct epithelial cells (cluster 8) **(Supplemental Figure S3B)**, both confirmed by quantitation of sample composition of each cluster **(Supplemental Figure S3C)**. Despite these findings, estimated nephron number was unaffected **(Supplemental Figure S2, C and D)**, consistent with the finding that *Cited1* is dispensable in kidney development (Boyle et al., 2007). We then analyzed the transcriptome by pseudobulking of each identified cluster with a specific focus on genes that are downregulated in mutant cells to identify the developmental programs that are disrupted or lost due to *Ilk* deficiency. An unbiased over-representation analysis (ORA) of down-regulated genes demonstrated that the top GO Biological Process terms were dominated by broad cellular metabolic programs almost ubiquitously **(Supplemental Figure S4, Supplemental Data 1)**.

**Figure 3:**
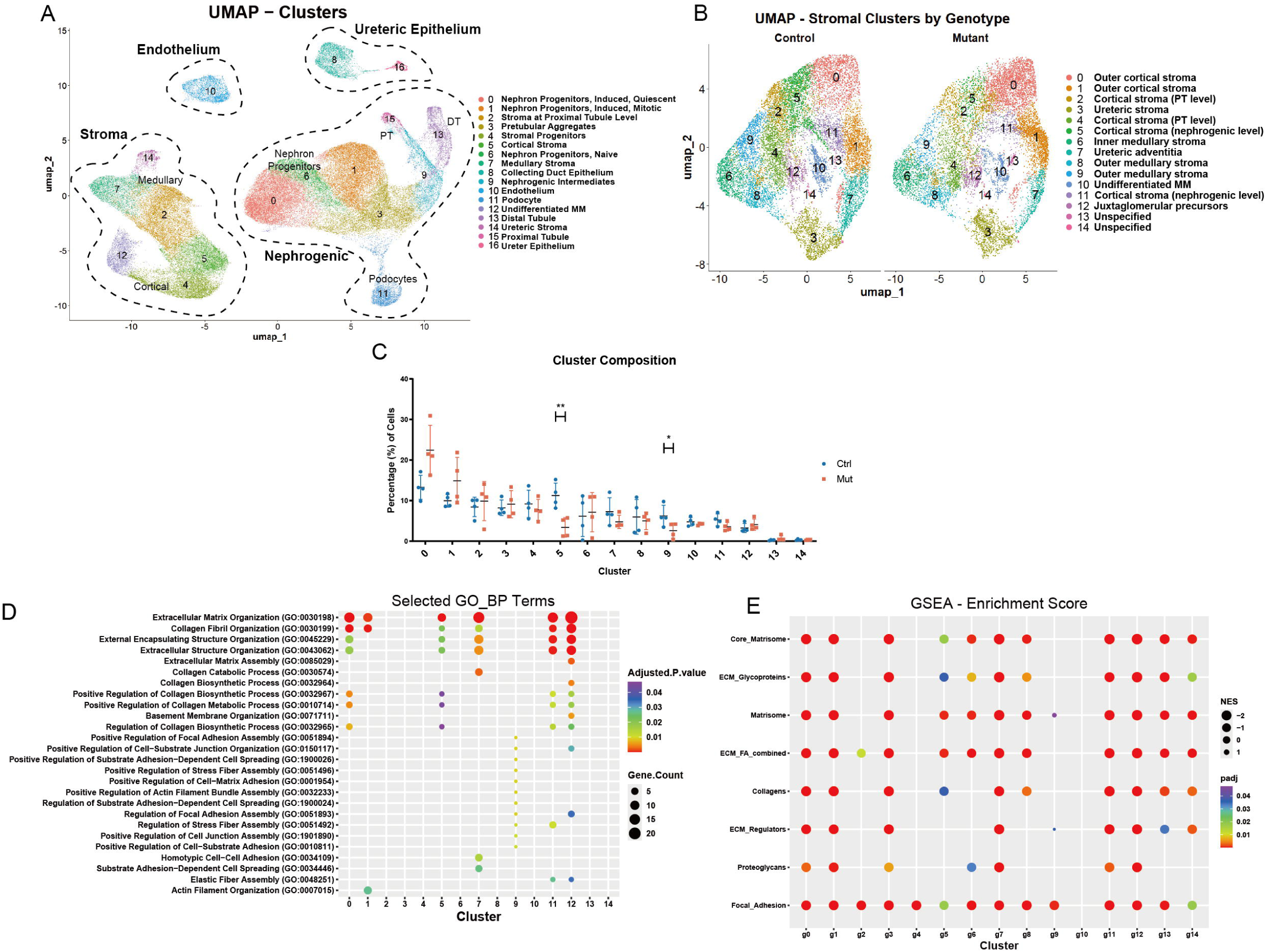
(A) UMAP reduction plot of single cell from whole E16.5 kidneys. 17 cell clusters were identified. (B) UMAP reduction of stromal subsets focusing on heterogeneity within stromal tissue. Re-cluster of stromal cells at a higher resolution identified 15 stromal cell clusters. (C) Cluster composition of each sample. Data are mean SD. * *p* < 0.05; ** *p* < 0.01 (empirical Bayes moderated *t*-test). (D) An unbiased over-representation analysis on genes downregulated in mutant kidneys for each pseudobulked stromal cell cluster identified gene ontology (biological process) terms related to extracellular matrix, cytoskeleton, and cell junction. (E) A Gene Set Enrichment Analysis targeting curated matrisome and focal adhesion gene sets confirmed downregulation of all tested gene sets with regional variation in stromal cell clusters.

To dissect the lineage and pathway-specific programs in greater detail, we subsetted the stromal cells which separate cleanly from other lineages on UMAP. Stromal populations were then clustered at a higher resolution, identifying 15 cell clusters (**Figure 3B**). These stromal clusters were annotated by the expression pattern of published E18.5 stromal heterogeneity markers (England et al., 2020) and then mapped to a published E16.5 kidney spatial RNA sequencing dataset (Jiang et al., 2025) to confirm their identities. Our data confirmed that the two independent annotation methods largely agree (**Figure 3B, Supplemental Figure S5, A and B)**. Quantitation of each stromal cell cluster confirmed a loss of cells in the phenotypic outer medullary region (stromal cluster 9) (**Figure 3C**). We again performed an unbiased ORA on pseudobulked stromal cell clusters focusing on downregulated genes **(Supplemental Figure S5C)**, and identified a change in extracellular matrix, focal adhesion, and cytoskeletal programs in many stromal clusters (**Figure 3D**). Judging by adjusted *p* values and gene counts, the extent of transcriptional alterations of the ECM and FA programs followed a corticomedullary gradient, with the effect being milder at the medullary end. Specifically, the phenotypic outer medullary cluster (stromal cluster 9) only demonstrated a mild downregulation in focal adhesion programs with no change in ECM components (**Figure 3D**). To rule out the effect of selected thresholds for fold-change and *p* values, we also interrogated the change in a threshold-free manner by gene set enrichment analysis (GSEA) on all detected genes ranked by the Wald statistic. We tested enrichment of a list of nested gene sets: the curated NABA_MATRISOME (Naba et al., 2012) and KEGG_FOCAL_ADHESION gene sets (Liberzon et al. 2011), two subcategories of the matrisome set (core matrisome and ECM regulators), and three subcategories of the core matrisome set (glycoproteins, collagens, proteoglycan). GSEA confirms the corticomedullary trend observed in ORA, and again demonstrates that the phenotypic cluster (stroma cluster 9) only carries FA program alteration at the transcriptional level (**Figure 3E**), with a small upregulation in matrisome and ECM regulators (**Figure 3E**) that is determined to be noise due to its sensitivity to ranking metrics. Overall, these data suggest that *Ilk* deletion generally disrupts ECM and FA programs in embryonic renal stroma, which is consistent with the well-known role of *Ilk*. However, different stromal populations react to *Ilk* deletion to a different extent. The population with strongest tissue-level phenotype demonstrated weak transcriptional alteration, which motivated investigation of non-transcriptional alterations in this region.

### Fibronectin expression is disrupted before tissue malformation in stromal *Ilk*-deficient kidneys

To identify FA- and ECM-proteins, whose corresponding transcripts are components of the transcriptional terms we identified by scRNASeq **(Supplemental Data 2)** and which are known to function with ILK, we performed immunostaining on kidney sections from control and mutant mice. Most FA components we tested did not demonstrate a change at E16.5 in the phenotypic outer medullary stroma except vinculin **(Supplemental Figure S6A)**. ECM components such as Netrin-1 and Laminin demonstrated no change at E16.5 **(Supplemental Figure S6C)**. Similarly, expression of Integrin α5 and β1, which are known to be dysregulated in previous *Ilk* knock-out models (Huveneers et al., 2008; Elad et al., 2013), was unchanged **(Supplemental Figure S6D)**. In contrast, other ECM components were altered. While decreased expression of Collagen VIII α2 in cortical stroma at E16.5 **(Supplemental Figure S6B)** and of dystroglycan in cortical, proximal tubule-level, and inner medullary stroma at E15.5 **(Supplemental Figure S6E)** was limited to non-phenotypic areas, and, therefore, not further investigated, one ECM component, fibronectin, was found to be altered in the outer medullary stroma.

While fibronectin appeared normal in cortical, proximal tubule-level, and inner medullary stroma (**Figure 4A**), a loss of fibronectin was observed in the outer medullary stroma at E16.5 (**Figure 4B**). We then examined fibronectin on E15.5 kidney sections, asking whether the change precedes the onset of the tissue-level phenotype. Our data confirmed a loss of fibronectin in the outer medullary stroma at E15.5 (**Figure 4B**) with no change in other regions **(Supplemental Figure S6F)**, placing fibronectin upstream of stromal tissue disruption temporally, and potentially mechanistically. Indeed, this possibility is supported by findings in the literature that fibronectin plays a critical role in initiation of ECM assembly (Garrison & Schwarzbauer, 2021), making it a potential primary target upstream of other histological and molecular alterations in *Ilk*-deficient stroma.

**Figure 4:**
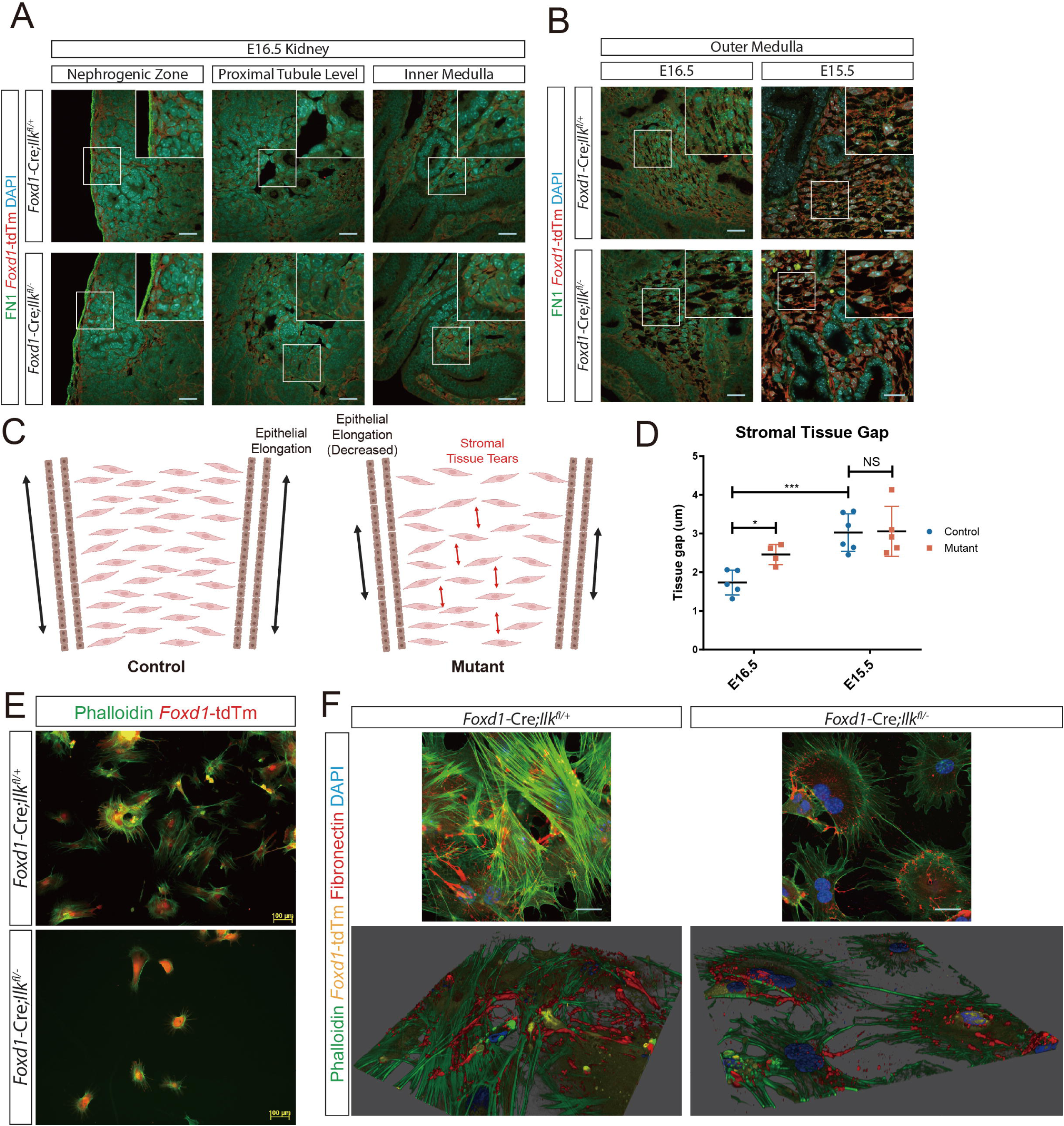
(A) Immunostaining of fibronectin in E16.5 kidneys. Images were captured along the corticomedullary axis. No difference was observed between control and mutant kidneys in nephrogenic zone, proximal tubule level, and inner medulla. Scale bar: 25 μm. (B) Immunostaining of fibronectin at E16.5 and E15.5. Images were captured at the outer medullary stroma. (C) A schematic demonstration of tissue tears characterized by increased gaps between adjacent stromal cells. Illustration is created at Biorender.com. (D) Quantitation of tissue gaps in control and mutant kidneys at E15.5 and E16.5. A difference between control and mutant kidneys is only detectable at E16.5. Data are mean SD. * *p* < 0.05; *** *p* < 0.001 (two-way ANOVA test). (E) Phalloidin staining of primary E15.5 stromal cells demonstrates disruptions in cytoskeleton organization and cell morphology. (F) Confocal images of primary E15.5 stromal cells immunostained to show fibronectin and phalloidin. 3D rendering of Z-stack images (bottom panels) demonstrates impaired fibronectin fibril assembly which normally occurs along actin filaments.

As we examined the tissue space anomaly at a higher resolution, we also noticed that the gaps between stripes of stromal cells, which are normally organized in a “rungs-on-a-ladder” pattern (Fusco et al., 2025) in the corticomedullary axis, were wider in mutant tissue than controls (**Figure 4B**). An illustration of the tissue pattern is shown in **Figure 4C**. This is suggestive of stromal tissue losing integrity and being torn open in the corticomedullary axis, an observation that has been previously made in nonrenal tissues when ECM anomalies occur (Marini et al., 2017; Neptune et al., 2003). We then quantified this change and confirmed that it was detectable starting at E16.5 (**Figure 4D**), coinciding with a shortened medulla (**Figure 2E**). These observations support a model in which mutant stroma and nearby genetically normal tissues create a mismatch in tissue expansion, causing tears in stroma and restricted elongation of nearby parenchyma.

A loss of fibronectin in tissue can result from various anomalies, including decreased transcription, disrupted translation and protein modification, disrupted secretion, or disrupted fibril assembly. To further interrogate these possibilities, we established primary stromal cell culture such that control and mutant cells can be placed in an identical condition for comparison, supplemented with sufficient soluble fibronectin molecules by serum additive in culture media. E15.5 stromal cells were isolated by FAC sorting for *Foxd1-*tdTomato fluorescent signal and seeded on culture plates. Unlike control stromal cells that can attach to plastic, mutant cells were found to be less adhesive and required surface treatment by poly-D-Lysine to attach to plastic. Phalloidin staining revealed abnormal cytoskeleton organization in mutant cells (**Figure 4E**), similar to *Ilk*-null fibroblast cell lines (Huveneers et al., 2008; Stanchi et al., 2009; Elad et al., 2013). We then examined fibronectin expression and assembly by immunostaining the primary cells. While control cells expressed fibronectin as thick fibrils, mutant cells demonstrated scattered fibronectin signals throughout the cell (**Figure 4F**). A 3-D rendering of Z-stack confocal images confirmed that fibronectin fibrils are organized along actin filaments in control cells, but not in mutant cells (**Figure 4F**). This is consistent with the fact that fibronectin fibrillogenesis requires cell contractility (Dzamba et al., 2009), which relies on the function of cytoskeleton. In summary, these data confirm that the loss of fibronectin in mutant stromal tissue results from anomalies intrinsic to mutant stromal cells.

### Embryonic renal stroma gains FA and ECM programs at transcriptional and non-transcriptional levels during the E15.5-E16.5 window

When comparing tissue-level phenotypes between control and mutant kidneys at E15.5-E16.5 of development, we observed that mutant E16.5 kidneys, when compared to E16.5 and E15.5 controls, demonstrate an intermediate state between E15.5 and E16.5 controls (**Figure 2E**, **Figure 4, B and D)**. This observation suggests a potential developmental arrest. We hypothesized that ECM and FA are critical developmental programs normally established in the stroma during this vulnerable window, which, when blocked by a loss of *Ilk*, result in disrupted tissue development.

To further investigate this hypothesis, we combined our single-cell transcriptomic data with published E15.5 control kidney single-cell datasets (Lawlor et al., 2019; Naganuma et al., 2021) and interrogated the developmental programs which are gained during this time window. E15.5 stromal cells were annotated using the same method as E16.5 stromal cells **(Supplemental Figure S7, A and B)**. Since clustering is driven by principal components that capture dataset-specific variance, and are, therefore, not directly comparable across analyses, we mapped all datasets onto the common spatial reference (Jiang et al., 2025) and labelled each cell based on its predicted location. This provided a shared, anatomically defined framework across developmental stages and genotypes. Comparing E15.5 and E16.5 control cells, an ORA on each pseudobulked region demonstrated a global developmental gain in matrix-related programs (**Figure 5A**), coinciding with the programs that were lost in E16.5 mutants compared to age-matched controls (**Figure 3, D and E; Figure 4B**). Interestingly, the phenotypic outer medullary region that demonstrated tissue-level ECM disruption without transcriptional alteration in the control-to-mutant comparison (**Figure 4B**) also gained these programs transcriptionally during normal development (**Figure 5A**). This is confirmed by a threshold-free GSEA (**Figure 5B**), which demonstrated a developmental gain in the outer medullary region in gene sets including all three subcategories of core matrisome (glycoproteins, collagens, and proteoglycans), ECM regulators, and FA during the E15.5 and E16.5 window (**Figure 5B**). These data suggest that the phenotypic outer medullary stroma indeed normally gains the developmental programs that *Ilk* deletion disrupts at a non-transcriptional level in this region, at the exact same time period, supporting the supposition that the observed tissue-level phenotypes represent a developmental arrest.

**Figure 5:**
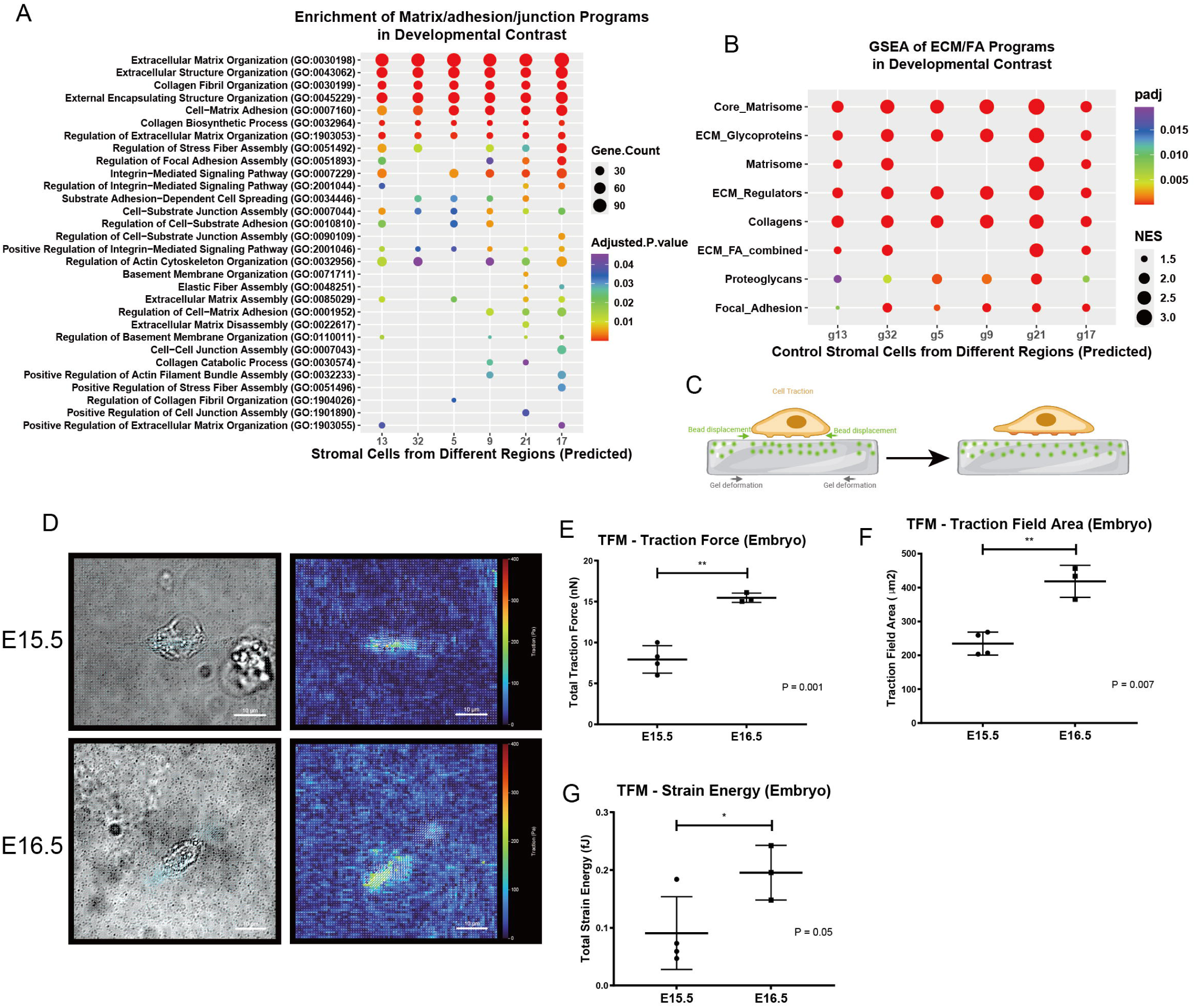
(A) An unbiased over-representation analysis comparing E15.5 and E16.5 wildtype stromal cells for each pseudobulked stromal region. Data demonstrate a global gain in extracellular matrix, cytoskeleton, and cell junction programs during development. (B) A Gene Set Enrichment Analysis targeting curated matrisome and focal adhesion gene sets confirmed a global developmental gain in all stromal regions. (C) A schematic demonstration of traction force microscopy assay. Illustration is created at Biorender.com. (D) Brightfield images (left panels) of primary stromal cells and heatmaps of traction field (right panels). (E-G) Quantitation of total traction force, traction field area, and total strain energy generated by E15.5 and E16.5 primary stromal cells. Data are mean SD. * *p* < 0.05; ** *p* < 0.01 (Student’s *t*-test).

Since our data indicates that the mutant phenotype emerges precisely at the E15.5-E16.5 window, if the mutant phenotype indeed reflects blockage of developmental programs, one would expect to see an abrupt change in these programs concurrently during normal development. We tested this by scoring matrisome and FA programs in pseudobulked stromal pixels from different regions in the spatial RNA sequencing dataset (Jiang et al., 2025) on E13.5, E15.5, E16.5, and E18.5 sections. While many regions demonstrated wide variation and inconsistent trends across replicate sections from different spatial resolutions, stromal pixels from two regions, namely, the outer medulla and middle medulla, demonstrated a continuous increase in the combined matrisome and FA program **(Supplemental Figure S7C, upper panels)**. Interestingly, the increase in FA- and matrisome-specific program scores demonstrated an abrupt deceleration in FA and an abrupt acceleration in matrisome **(Supplemental Figure S7C, middle and lower panels)**, both occurring at the time of the E15.5-E16.5 window. While the spatial dataset does not contain sufficient replicates measured in a consistent method to be conclusive, this finding underscores the importance of this window in stromal development and supports the concept that the onset of the mutant phenotype is related to the normal developmental timeline.

Since the phenotype in mutant kidney outer medullary stroma was observed at a non-transcriptional level (**Figure 4B**), we then investigated the presence of a non-transcriptional developmental change that normally occurs at this critical window, which could confer the developing stroma susceptible to disruptions by *Ilk* deficiency at this stage. Since fibronectin is the most likely primary target, we pursued cell traction, which is known to be required for fibronectin assembly (Dzamba et al., 2009) and is known to depend on normal cytoskeleton and focal adhesion (Wang, 2010), both of which were disrupted in *Ilk*-deficient cells (**Figure 3E**; **Figure 4F**). We performed traction force microscopy (TFM) on E15.5 and E16.5 stromal cells, which assesses cell traction by seeding cells on an ECM-crosslinked substrate with embedded fluorescent beads and measuring bead displacement (**Figure 5C).** Our data indicate that E16.5 cells generate a significantly higher total traction force and total strain energy than E15.5 cells, accompanied by a larger traction field (**Figure 5, D-G)**, demonstrating that stromal cells undergo a cell behavior-level developmental change during this critical window.

Together, these data demonstrate that critical developmental programs related to FA function and ECM assembly are established in embryonic renal stroma at both transcriptional and non-transcriptional levels, both occurring during the E15.5-E16.5 time window. While these programs are ubiquitously gained by all stromal populations at multiple levels during development, our data suggest that *Ilk* regulates these programs at different levels in different stromal populations, and when deleted, results in tissue phenotypes that impact specific cell populations.

## Discussion

In this study, we investigated the regulatory role of ILK during development of embryonic renal stroma in a mouse model with *Ilk* deletion specific to renal stromal progenitors. Our results demonstrate a shortened renal medulla and disruption of ECM accompanied by progressive loss of tissue integrity and formation of tissue tears, which eventually developed into fluid-filled tissue spaces in the medullary interstitium followed by stromal cell death. Using single-cell RNA sequencing, we identified ECM and FA programs as targets downstream of *Ilk* and established differential response to a loss of *Ilk* across heterogenous stromal cell populations, at transcriptional and non-transcriptional levels. Using public time course single-cell and spatial RNA sequencing datasets, we demonstrated a global gain of ECM and FA programs, the same developmental programs that were blocked when *Ilk* is deleted, in renal stroma during development. Combining these data with our histological observations, we identified a key developmental window, E15.5-E16.5, for establishment of stromal function. A model summarizing the developmental process and disruptions by the mutant is provided in **Figure 6**.

**Figure 6:**
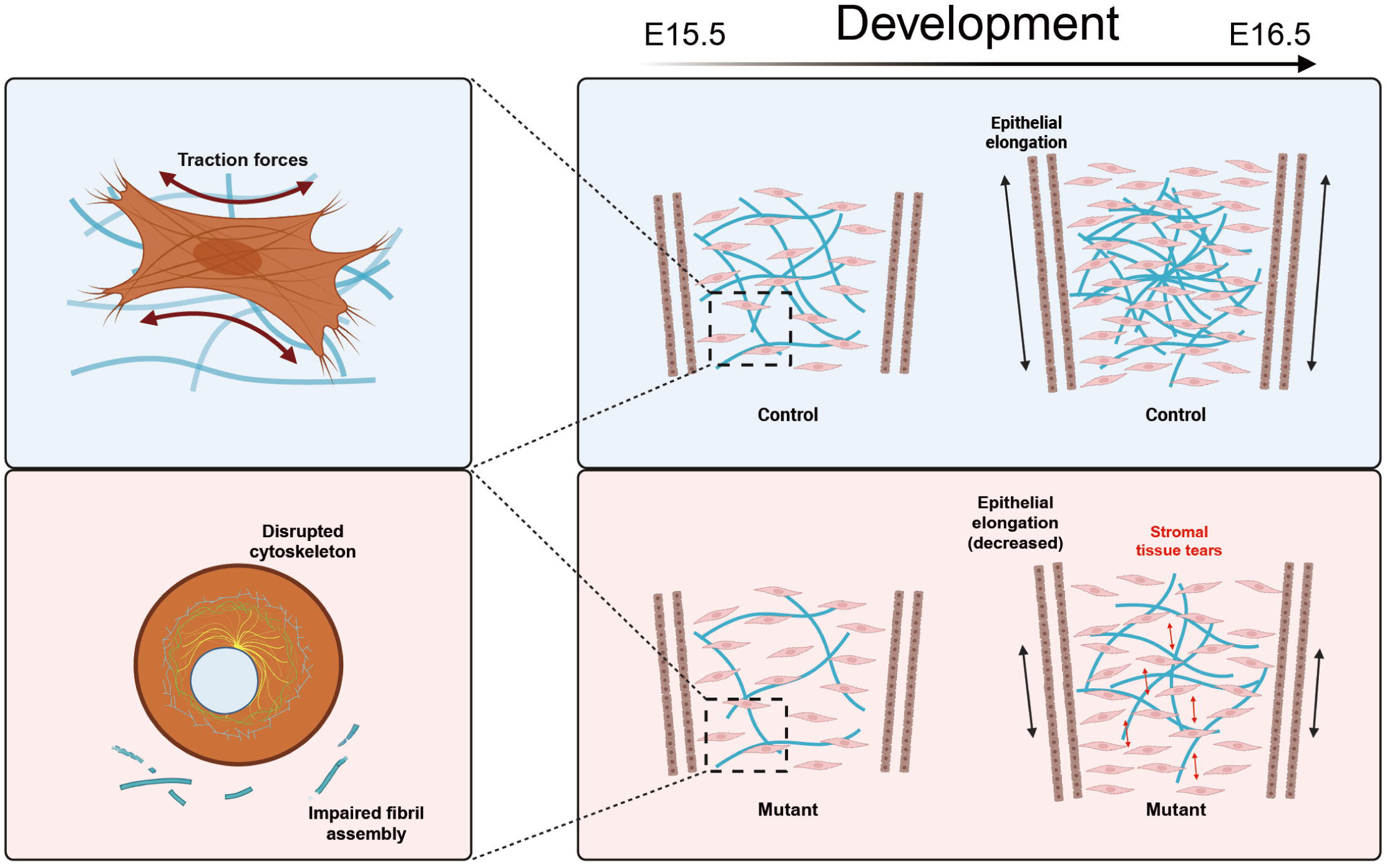
A schematic demonstration of findings in this study. Normally, the renal stroma gains expression of ECM and FA programs at cellular and tissue levels both transcriptionally and non-transcriptionally during a critical developmental time window, E15.5-E16.5. These programs can be disrupted by a loss of *Ilk* specific to the stroma, causing tissue tears in the corticomedullary axis and loss of cellularity in the outer medullary stroma, arising precisely at the same time window, accompanied by loss of fibronectin fibrils, and affecting elongation of epithelial lineages as well. Out data demonstrate that loss of fibronectin and cellular anomalies precede tissue-level disruptions, suggesting a model in which cellular activities, regulated by *Ilk*, control tissue integrity through ECM organization. Illustration is created at Biorender.com.

In *Ilk*-null mutants, we observed a fluid-filled medullary region in developing kidneys, resembling hydronephrosis. Although this phenotype might initially suggest a primary defect located in the urinary collecting system (that is, the urinary pelvis and/or ureter), our data show that it originates in the stromal lineage that carries the mutation and develops as fluid-filled tissue spaces in the medullary interstitium. The source of the fluid is unknown. Our gross anatomical and histologic observations suggest that the tissue spaces are not connected to the urinary tract. Therefore, it is not likely that urine directly entered the tissue spaces through a tissue opening. However, leakage of epithelial tubules cannot be ruled out. In mouse embryonic kidneys, urine production and filtration starts as early as E16.5 (Rasouly & Lu, 2013), coinciding with the onset of tissue disruptions. Since no anomaly was detected by single-cell RNA sequencing and immunostaining in the epithelial compartments of the medulla at E16.5, epithelial leakage, if present, likely arises at a later stage when stromal tissue tears transformed into tissue spaces, downstream of anomalies in the stromal compartment.

Our data suggest that death of stromal cells is a consequence of ECM and tissue disruptions in *Ilk*-null stroma, supported by the following temporal sequence: ECM disruption at E15.5 preceding all visible disruptions, followed by loss of cellularity and formation of tissue tears at E16.5 without cell death; followed by tissue tears evolving into tissue spaces at E17.5-E18.5, and followed by cell death eventually observed at P5. A possible mechanism underlying cell death that fits this sequence is anoikis, a special form of cell death that occurs when cells detach from the surrounding ECM (Frisch & Francis, 1994). While it is certain that anoikis is downstream of ECM disruptions, at least temporally, where anoikis sits in the sequence of tissue-level anomalies is unknown. It is possible that the ECM anomaly in mutant tissue directly triggers the anoikis program, which in turn, contributes to formation of tissue spaces. Meanwhile, the loss of stromal cellularity and the tissue tears could also cause a sparsely populated environment that mimics detachment, which in turn, triggers anoikis. Once the tissue-level phenotype emerges, loss of cellularity, ECM anomaly, compromised tissue integrity, and anoikis can together form a self-reinforcing feedback loop. Time course observation at a very high temporal resolution may help to resolve the exact sequence of these events, but the fact that so many progressive events occur in a small window that only lasts for a few days makes this challenging. As we mainly focused on the primary anomalies at earlier stages, our single-cell data did not provide an answer to this question.

The initial cause of the loss of cellularity in the phenotypic area is still unknown. As cell death and proliferation have been ruled out, we have considered cell migration as a possible cause: ECM is required for cell migration, and our data demonstrate a disruption in ECM assembly in mutant kidneys. However, to conclude that disrupted cell migration causes loss of medullary stromal cells, one must first establish that normal stroma cells migrate into the medulla during embryonic kidney development. Although the stromal progenitor cells are located in the cortical region, whereas the differentiated stromal cells are located in the medulla, such relocation can be explained by distributed tissue expansion or progenitor cells giving rise to new cells and pushing new cells toward the medulla, rather than cells actively migrating relative to surrounding cells, which is the defining feature of cell migration. Distinguishing these possibilities will require advanced live tissue imaging techniques and reporters that sparsely label individual stromal cells and their neighbors rather than an entire lineage.

Our data demonstrate a shortened medulla accompanied by stromal tears in the corticomedullary axis. This phenotype could reflect a mechanical tension at the tissue interface: as stromal expansion fails to match the extension of epithelial tubules, the extending epithelial tubules can exert traction to the abnormal stroma, causing the stroma to tear, while the stroma in turn limits epithelial extension. The biomechanical explanation contains three subtly different scenarios: (1) increased radial tension from epithelial tissues alone caused the stroma to tear; (2) the stroma is less resistant to tissue tension due to a loss of cells and abnormal ECM, while the tension is unchanged; (3) a combination of both (1) and (2). As ECM is critical to maintenance of tissue integrity, it is very unlikely that tissue toughness remains unaffected, hence the combination is most likely the case. These possibilities can be distinguished by various biomechanical analyses, such as stiffness measurement by atomic force microscopy and shear wave elastography, and tension measurement by cell shape analysis (Roffay et al., 2021) and injected force sensors (Lee et al., 2019). Alternatively, the tissue tearing and shortening may not be purely mechanical, as molecular regulations such as the planar cell polarity pathway could also explain a change in directional cell division and tissue extension.

*Ilk* has previously been studied at a cellular level, demonstrating an essential role in FA maturation (Stanchi et al., 2009). Specifically, *Ilk* has demonstrated capacity to regulate fibronectin fibrillogenesis via integrin α5β1 in a cultured fibroblast cell line (Huveneers et al., 2008; Elad et al., 2013). As well, fibronectin is known to be a foundational component in ECM assembly that other components depend on to incorporate into the ECM (Garrison & Schwarzbauer, 2021). Consistent with these *in vitro* findings, our data also demonstrate that in *Ilk*-deficient stroma, disruption of fibronectin is temporally upstream of all other tissue-level anomalies. Our data demonstrate the consequences of disruptions in *Ilk* and fibronectin function at the tissue level. Moreover, results, reported here, reveal that the ECM anomaly downstream of *Ilk* deficiency in stromal tissue differs from region to region. Similarly, *Ilk* deletion results in various phenotypes in other renal lineages: loss of contact inhibition and collecting duct obstruction in the ureteric lineage (Smeeton et al., 2010), and alterations in glomerular basement membrane, foot process effacement, and loss of slit diaphragm in podocytes. Difference in phenotype in different tissue or organs is also seen with other ECM molecules, such as *COL1A1* and *COL1A2* mutations causing skin fragility, muscle weakness, and dentinogenesis imperfecta (Marini et al., 2017), and *FBN1* mutations causing Marfan syndrome, characterized by aortic root aneurysm and dissection, lens dislocation, long limbs, and lung emphysema (Neptune et al., 2003). These varied phenotypes reflect a wide variation in ECM architecture and requirement for *Ilk* and FA function as a result of different mechanical demands, signaling cues, and resident cell behavior.

Apart from the phenotypic outer medullary stromal cluster (stromal cluster 9) that demonstrated reduced cells in mutant kidneys, our data also suggests a loss of cells in the inner cortical stroma (stromal cluster 5) (**Figure 3C**). We did not further pursue this population, since stromal cells in this region appeared normal on tissue sections. We observed a mild increase in a nearby population on UMAP that represents outer cortical stroma (stromal cluster 0) (**Figure 3B**), although the change in this population did not reach statistical significance. This change in cortical cell proportion likely reflects a change in transcriptomic signature rather than an actual loss of cells, resulting in some cells from inner cortical stroma being categorized as outer cortical.

ECM-related programs have been implicated in renal stromal development. Spatial RNA sequencing on embryonic kidney section in timeseries has reported an increase in ECM-related ligand-receptor pairs comparing early (E11.5-E13.5) and late (E15.5-E18.5) embryonic sections (Jiang et al., 2025). However, this was investigated at a relatively coarse temporal resolution. Our data indicates that significant developmental changes can occur within a 24-hour window, and such changes are likely masked when stages are pooled. In particular, our data identify E15.5-E16.5 as a critical window for establishment of these developmental programs. Our analyses highlight the importance of this narrow window.

## Methods

### Mouse models

To generate mice in which *Ilk* deficiency is generated specifically in the *Foxd1*+ stromal lineage, *Foxd1*-*eGFP/Cre* mice (Humphreys et al., 2008) (referred to as *Foxd1^Cre^*) were first crossed with *Ilk^+/-^* mice generated by NLS-Cre, resulting in *Foxd1^Cre/+^*;*Ilk^+/-^*offsprings. Mice homozygous for floxed *Ilk* (Terpstra et al., 2003) alleles (referred to as *Ilk^fl/fl^*) were crossed with *Rosa26-CAG-LSL-tdTomato* mice (referred to as *Rosa26^tdTm/tdTm^*) (purchased from Jackson Laboratories) in which expression of a red fluorescent protein is prevented by a floxed stop cassette and can be activated in the presence of Cre recombinase (Madisen et al., 2010). The offspring were intercrossed to maintain a homozygous strain (*Ilk^fl/fl^*;*Rosa26^tdTm/tdTm^*). *Foxd1^Cre/+^*;*Ilk^+/-^* male mice were then mated with *Ilk^fl/fl^*;*Rosa26^tdTm/tdTm^* female mice, resulting in *Foxd1^Cre/+^*;*Ilk^fl/-^;Rosa26^tdTm/-^*offspring with lineage-specific *Ilk* deletion by *Foxd1*-Cre in the descendant cells of the *Foxd1*+ stromal progenitors (referred to as stromal *Ilk*-deficient), which are labelled by red fluorescent signal. All animals were housed at The Centre for Phenogenomics (TCP; Toronto, Canada).

### Genomic DNA isolation and transgenic mice genotyping

To genotype mouse offsprings, tissue samples were collected from mouse embryo limb tissue or ear notches and tail clips from post-natal mice. DNA was extracted using the Advanced DNA Fast Extract Buffers Kit (Wisent). Alleles were amplified by PCR using the primers in **Supplemental Table S1** and distinguished by the size of PCR products using agarose gel electrophoresis. For *Foxd1^Cre/+^*;*Rosa26^tdTm/tdTm^* mice, expression of the *Foxd1^Cre^* allele was determined by observing red fluorescence signal with epifluorescence microscope (Zeiss).

### Culture of Primary Stromal Cells

FAC-sorted stromal cells were collected by centrifugation at 200 g for 5 minutes. Cells were counted with a Countess 3 automated cell counter (Invitrogen) and then seeded on 8-well plastic chamber slides (Falcon) at 6000 live cells per well in 0.5 mL of media. Slides were treated with poly-D-lysine (Sigma-Aldrich) at 0.1 mg/mL at 4°C overnight before stromal cells were seeded. Seeded cells were cultured in DMEM/F12 (GIBCO) media supplemented with 10% FBS (GIBCO) and 1% Penicillin Streptomycin (P/S) (GIBCO), in 5% CO_2_ at 37°C.

### Intrapelvic dye injection

Mouse kidneys were dissected at post-natal day 0 and day 5 (P0 and P5). Intrapelvic dye injection was performed as previously described (Airik et al., 2006). Bromophenol blue was injected into the renal pelvis of whole urogenital systems using pulled glass capillary needles.

### Tissue fixation and histology

Dissected kidneys were preserved in 4% PFA overnight at room temperature and stored in 70% ethanol at 4°C until processing. Kidneys were then embedded in paraffin and serial-sectioned at the sagittal plane exhaustively at 5 μm increments. For embryonic kidneys at embryonic day 14.5 (E14.5) or earlier, decapitated embryos were processed and embedded intact and later trimmed to demonstrate kidney tissue when generating tissue sections. Additionally, to ensure correct orientation of kidneys for sectioning at the sagittal plane, E15.5 and E16.5 kidneys were pre-embedded in a mixture of gelatin (Sigma-Aldrich) and low melting point agarose (Thermo Scientific) at final concentrations of 2% and 2.5% respectively (McClelland et al., 2015) prior to embedding in paraffin. Paraffin embedding, sectioning, H&E staining, PAS staining and TUNEL staining were performed by the Pathology Lab at The Centre for Phenogenomics (TCP; Toronto, Canada).

### Immunofluorescence

Kidney sections were deparaffinized in xylene and rehydrated through an ethanol gradient. Antigen retrieval was performed in sodium citrate buffer (pH 6.0) with a microwaveable pressure cooker. Sections were blocked with DAKO Serum-Free Protein Block (Agilent) for 1 hour at room temperature. Primary antibodies were incubated overnight at 4°C. Sections were then incubated in Alexa Fluor secondary antibodies (Thermo Scientific) diluted at 1:1000 and DAPI (Sigma-Aldrich) at a final concentration of 10 μg/mL, washed with PBS, and mounted with DAKO Fluorescence Mounting Medium (Agilent). Stained sections are imaged with an epifluorescence microscope (Zeiss), a confocal microscope (Nikon A1R or Leica Stellaris 8 FALCON), or a slide scanner (3DHistech Pannoramic).

### Quantification of nephron formation

E16.5 kidneys were serial-sectioned as described in **Tissue Fixation and Histology** and PAS-stained. Glomeruli were counted in each level of serial sections and cumulated throughout all levels to represent a kidney as previously described (Cain et al., 2009)

### Quantification of Development of the Renal Medulla

Embryonic kidneys were serial-sectioned as described in **Tissue fixation and histology**, For each kidney, a section reflecting the multiple collecting ducts lining in parallel to the corticomedullary axis and intersecting with each other at the papilla was considered the representative mid-sagittal section and was used to quantify the area of medulla. If no section showing such pattern was collected, two sections containing the largest area were considered to “flank” the mid-sagittal plane, and the average medullary area of these two sections was computed to represent the medullary area at the mid-sagittal plane. For each kidney, the representative section(s) was immunostained for LEF1 and counterstained for RFP and DAPI. Slides were imaged with an epifluorescence microscope (Zeiss). Images were stitched together to reconstruct the mid-sagittal plane of a whole kidney using Photoshop software (Adobe). Stitched images were processed with ImageJ software (The National Institutes of Health, United States) for measuring the areas of the medulla and the kidney section. The medulla area to kidney area ratio was then computed.

### Assessment of Tissue Tears in Outer Medullary Stroma

A mid-sagittal section was selected from each kidney and immunostained for RFP signal expressed in stromal cells. On each section, each outer medullary stromal region defined by two adjacent epithelial tubules that are at least three stromal cells apart was examined. The gaps between adjacent stromal cells along the corticomedullary axis were measured. For each region, at least 100 μm of stromal tissue along the corticomedullary axis and 20 cell gaps were measured, and the average gap distance for each region was computed. Results were then averaged across all measured regions on a section to represent a kidney.

### Library Preparation for Single-Cell RNA Sequencing of E16.5 Mouse Kidneys

Kidneys were dissected from mutant and control kidneys at E16.5, each group containing 4 biological replicates collected from two litters. To meet the cell number required for analysis, each biological replicate contains 2 pairs of kidneys from 2 littermate embryos. Kidneys were snap-frozen in liquid nitrogen and stored at −80°C until processed. Processing of kidney samples and RNA sequencing were performed at Princess Margaret Genomics Centre (PMGC; Princess Margaret Cancer Research Tower, Toronto, Canada). Single cells were isolated to perform single-cell RNA sequencing using a Chromium Fixed RNA Profiling Reagent Kit and Single Cell Gene Expression Flex Fixed RNA Profiling workflow (10x Genomics).

### Analysis of Single-Cell RNA Sequencing Datasets

Raw FASTQ reads were aligned to the mm10 genome and converted to gene-barcode matrices using Cell Ranger pipeline (10x Genomics) with recommended settings by the manufacturer. The data set was subsequently processed using a Seurat V5.0 (Hao et al., 2024) workflow. Cells with a mitochondrial gene percentage of greater than 8% and library size below 7600 genes per cell were excluded. Cell doublets were removed using DoubletFinder (McGinnis et al., 2019), and cells representing blood cells, platelets and macrophages were excluded. The cells were clustered based on shared nearest neighbor graphs using Seurat’s FindNeighbors and FindClusters functions. The cell composition of each cluster was computed per sample, and the proportion of each cluster in each sample was compared between control and mutant using logit-transformed proportions (Phipson et al., 2022) with a moderated *t*-test in limma (Ritchie et al., 2015). Pseudobulk analysis was performed on each cluster using the AggregateExpression function. Genes differentially expressed (DEGs) between control and mutant data sets were identified in post-pseudobulk data set using DESeq2 (Love et al. 2014) and screened with adjusted P < 0.05 and |log2FC| > 0.58. An unbiased overrepresentation analysis was performed on selected DEGs using ENRICHR (Chen et al., 2013; Kuleshov et al., 2016). A Gene Set Enrichment Analysis was performed with fgsea (Korotkevich et al., 2021) on genes ranked by the DESeq2 Wald statistic. Enrichment was tested by gene-set permutation. A competitive rank test (limma cameraPR; Wu & Smyth, 2012) was used as a confirmatory test. Each cell in our dataset was mapped to a public spatial transcriptomic dataset (Jiang et al., 2025) to predict its location using Seurat’s anchor-based label transfer. Program scores were computed with an expression-matched module score (Tirosh et al., 2016).

### Fluorescence-Activated Cell Sorting (FACS)

Dissected mutant and control kidneys were digested in 1 mg/mL Collagenase B (Roche) in Accumax (Cedarlane) at 37°C for 1.5 hours and then gently pipetted to dissociate. The dissociated tissue was centrifuged at 200 g for 5 minutes and the cell pellet was resuspended in PBS containing 1 mM EDTA and 1% FBS. The cell suspension was filtered through a 70 μm cell strainer (BD Falcon). FACS was performed at the SickKids-UHN Flow Cytometry Facility (Toronto, Canada). Stromal descendant cells were separated using a Sony MA900 Multi-Application Cell Sorter by expression of tdTomato signal in the stromal lineage.

### Traction Force Microscopy

Traction force microscopy assay was performed using a modified method from previous reports (Cretu et al., 2010; Liu et al., 2024). Glass cover slips (#1, 25 mm, Chemglass Life Sciences) were silanized as previously described (Cretu et al., 2010). Polyacrylamide gel mixture containing fluorescent beads (1 μm mean particle size, #L4655, Sigma-Aldrich) was prepared as described (Liu et al., 2024). 30 μL of gel and bead mixture was added on each cover slip to form a glass-gel-glass sandwich (Cretu et al., 2010). 3.5 mm round dishes with a separate cover slip bottom (ibidi) were used for cell culture. The original cover slip bottom of the dish was removed by heating with a heat block and replaced with the cover slip that holds the fabricated polyacrylamide gel. After fabrication, the surface of the gel was functionalized as described (Liu et al., 2024) using a 365nm UV LED nail lamp. Functionalized gel was then treated with 1mg/mL Matrigel (Corning) at 4°C overnight to crosslink with ECM components to support cell adhesion. E15.5 and E16.5 kidneys were dissociated as described (Liu et al., 2024) and filtered with a 70 μm cell strainer (BD Falcon). The cell suspension was then seeded on the prepared gel and cultured for 12 hours in DMEM/F12 (GIBCO) media supplemented with 10% FBS (GIBCO) and 1% Penicillin Streptomycin (P/S) (GIBCO), in 5% CO_2_ at 37°C. As previously described (Liu et al., 2024), during traction force measurement, cells were treated with 4% PFA to create a loose state comparing to the untreated full traction state and then stained for a stromal marker, PBX1, to confirm their stromal identity. A confocal microscope with stage coordinate registration (Nikon A1R) was used to image the cells and the beads at three stages: full traction, loose state, and stained state. A 3D printed scaffold was used to hold the fabricated culture dish such that dishes can be placed on the stage at the same location each time to reload registered cell coordinates. Images were processed and analyzed with MATLAB. Loose and tense images were aligned with a rigid-body transform estimated by FFT phase correlation using MATLAB’s Image Processing Toolbox. Bead displacements were measured by particle image velocimetry (PIV) (Willert & Gharib, 1991; Crocker & Grier, 1996; Westerweel et al., 1997; Westerweel & Scarano, 2005). Traction stresses were computed by Fourier-transform traction cytometry (FTTC; Butler et al., 2002; Sabass et al., 2008). Cells with a signal-to-noise ratio (SNR) lower than 2 were excluded from analysis.

## Quantitation and Statistical Analysis

Statistical analyses were performed using Microsoft Office Excel software and Prism. The differences in the proportion of medulla in kidney mid-sagittal area and in stromal tissue gaps were each tested by a two-way ANOVA test. The change in cell cluster composition in single-cell dataset was tested with an empirical Bayes moderated *t*-test. Enrichment of gene sets in GSEA was tested by a permutation test and a Wilcoxon–Mann–Whitney rank-sum test. The changes in traction field, total traction force, strain energy, and estimated nephron number were tested by a Student’s *t*-test. A *p* value (or adjusted *p* value where appropriate) of < 0.05 was considered statistically significant. Error bars denote standard deviation (SD).

## Author contributions

Xiangyue Hu: Conceptualization, Methodology, Investigation, Data curation, Formal analysis, Software, Validation, Visualization, Writing - original draft, Writing - review & editing.

Norman Rosenblum: Conceptualization, Methodology, Resources, Funding acquisition, Supervision, Project administration, Writing - review & editing.

All authors have read and accepted the data being presented in the manuscript.

## Acknowledgement

We thank Dr. Troy Ketela, Genevieve Mailhot, Mary Shi, and Christine Pham from the Princess Magaret Genomics Centre for their assistance with single-cell RNA sequencing; Dr. Kimberly Lau and Paul Paroutis from the Imaging Facility at the Hospital for Sick Children for technical support with imaging; the SickKids-UHN Flow Cytometry Facility for support with cell sorting; and the Pathology Lab at The Centre for Phenogenomics for histological sample preparation. We thank Drs. Alex Hughes and Jiageng Liu for generously sharing their protocol and helping us optimize our traction force microscopy assay. We thank Zihao Chen for co-designing and 3D-printing a sample holder for cell coordinate registration and reloading. We thank Yun-kyo Kim, Dr. Min Zhu, Dr. Sevan Hopyan, Dr. Sara Qubisi, Dr. Jon Lacanlale, and Dina Greenberg for helpful discussions throughout this project. Finally, we thank Rukhsar Sultana and Nalma Uy for providing administrative support that allowed this project to be completed, under considerable constraints.

## Data availability

The RNA-sequencing data discussed in this publication will be accessible through GEO Series accession number GSEXXX.

