## Supplemental Materials for "Integrin-Linked Kinase Controls Tissue Integrity in Embryonic Renal Stroma Via Expression of Fibronectin During a Critical Developmental Time Window"

**Primers used for genotyping**

| *Foxd1*-Cre: 10559 | GGC ATT AAA GCA GCG TAT CC |
| --- | --- |
| *Foxd1*-Cre: 11122 | CTG TTC CTG TAC GGC ATG G |
| *Foxd1*-Cre: oIMR9535 | GGC ATT AAA GCA GCG TAT CC |
| tdTomato: oIMR9020 | AAG GGA GCT GCA GTG GAG TA |
| tdTomato: oIMR9021 | CCG AAA ATC TGT GGG AAG TC |
| tdTomato: oIMR9103 | GGC ATT AAA GCA GCG TAT CC |
| tdTomato: oIMR9105 | CTG TTC CTG TAC GGC ATG G |
| *Ilk* del Fwd | CCA GGT GGC AGA GGT AAG TA |
| *Ilk* del Rev | CAA GGA ATA AGG TGA GCT TCA GAA |
| *Ilk* flox/WT Fwd | CAA GGA ATA AGG TGA GCT TCA GAA |
| *Ilk* flox/WT Rev | AAG GTG CTG AAG GTT CGA GA |

Supplemental Table S1. Primers used for mouse offspring genotyping.

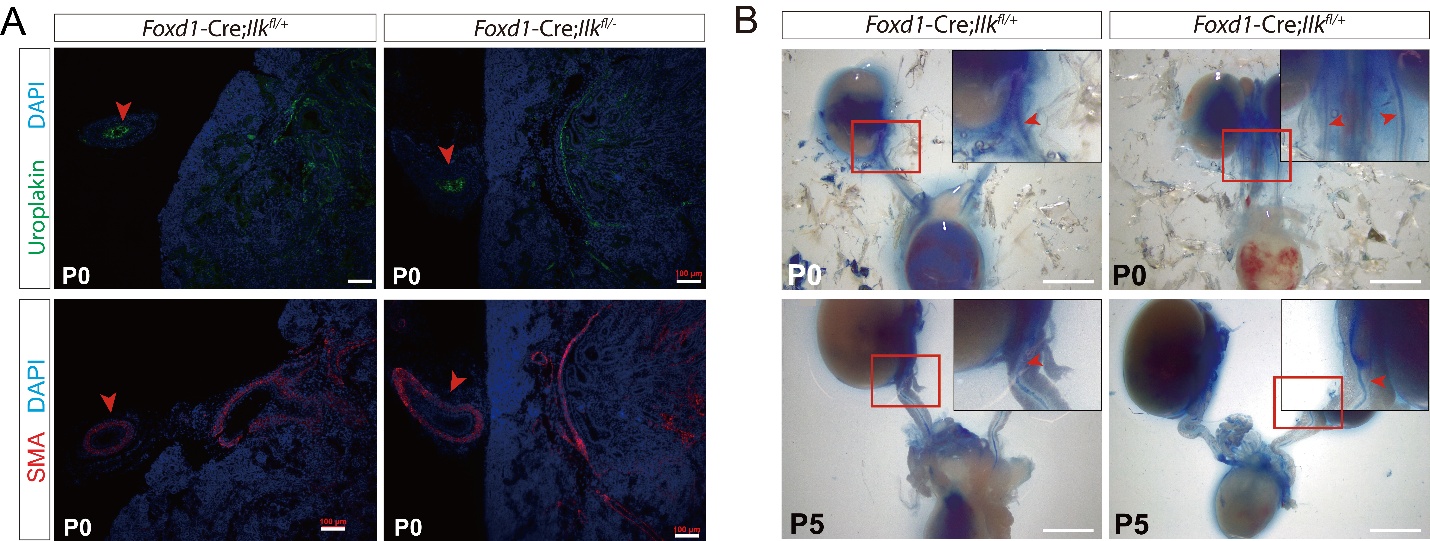

Supplemental Figure S1

(A) Immunostaining of urothelium and ureteric smooth muscle. Scale bars: 100 μm (B) Intrapelvic dye injection shows no obstruction at the ureteropelvic junction. Scale bars: 2mm.

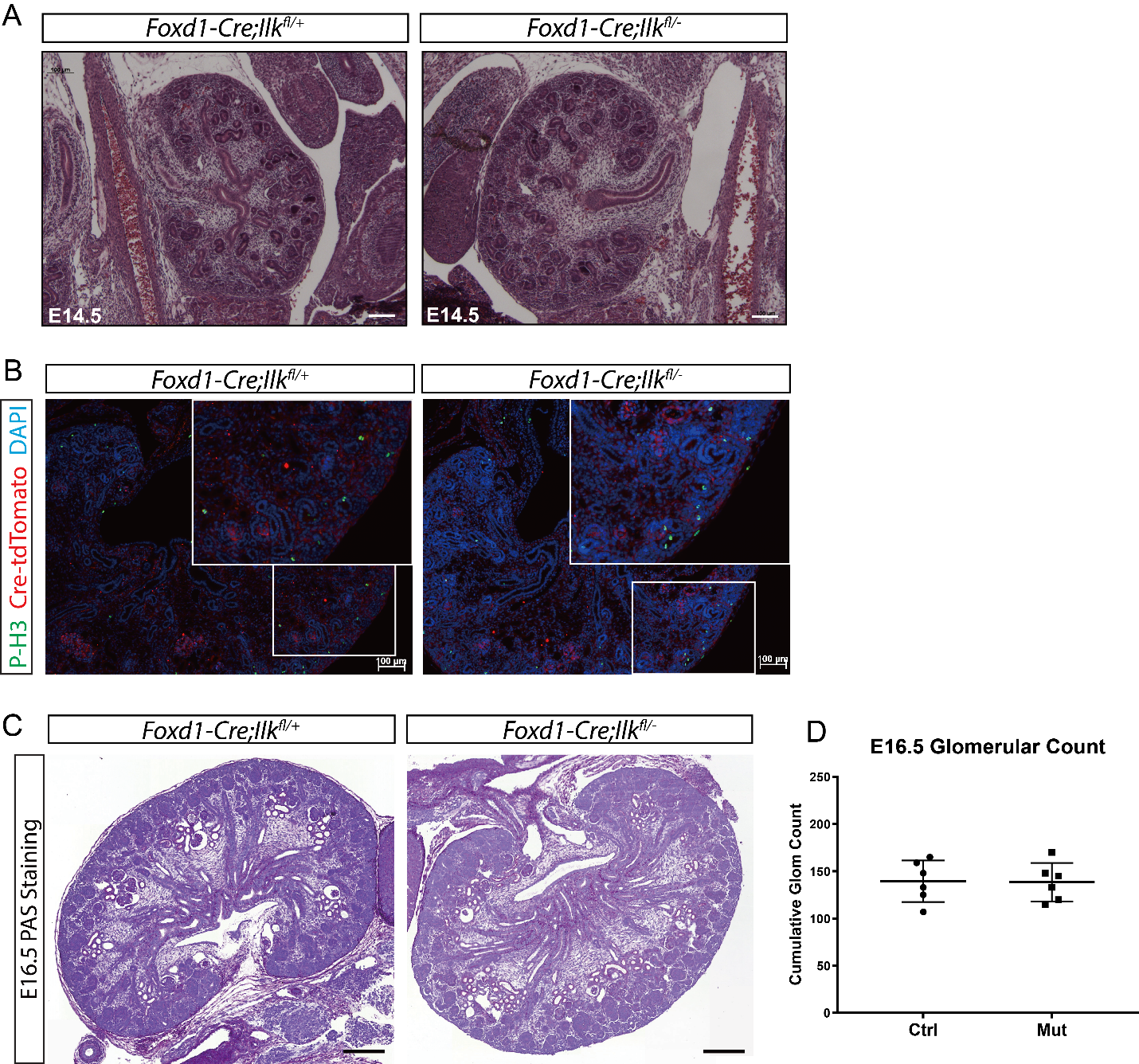

Supplemental Figure S2

(A) H&E staining of E14.5 kidneys. Scale bars: 100 μm. (B) Immunostaining of phosphor-Histone H3 on E16.5 kidney sections. Scale bars: 100 μm. (C) PAS staining of E16.5 kidney mid-sagittal sections. Scale bars: 200 μm. (D) Quantitation of glomerular numbers in E16.5 kidneys.

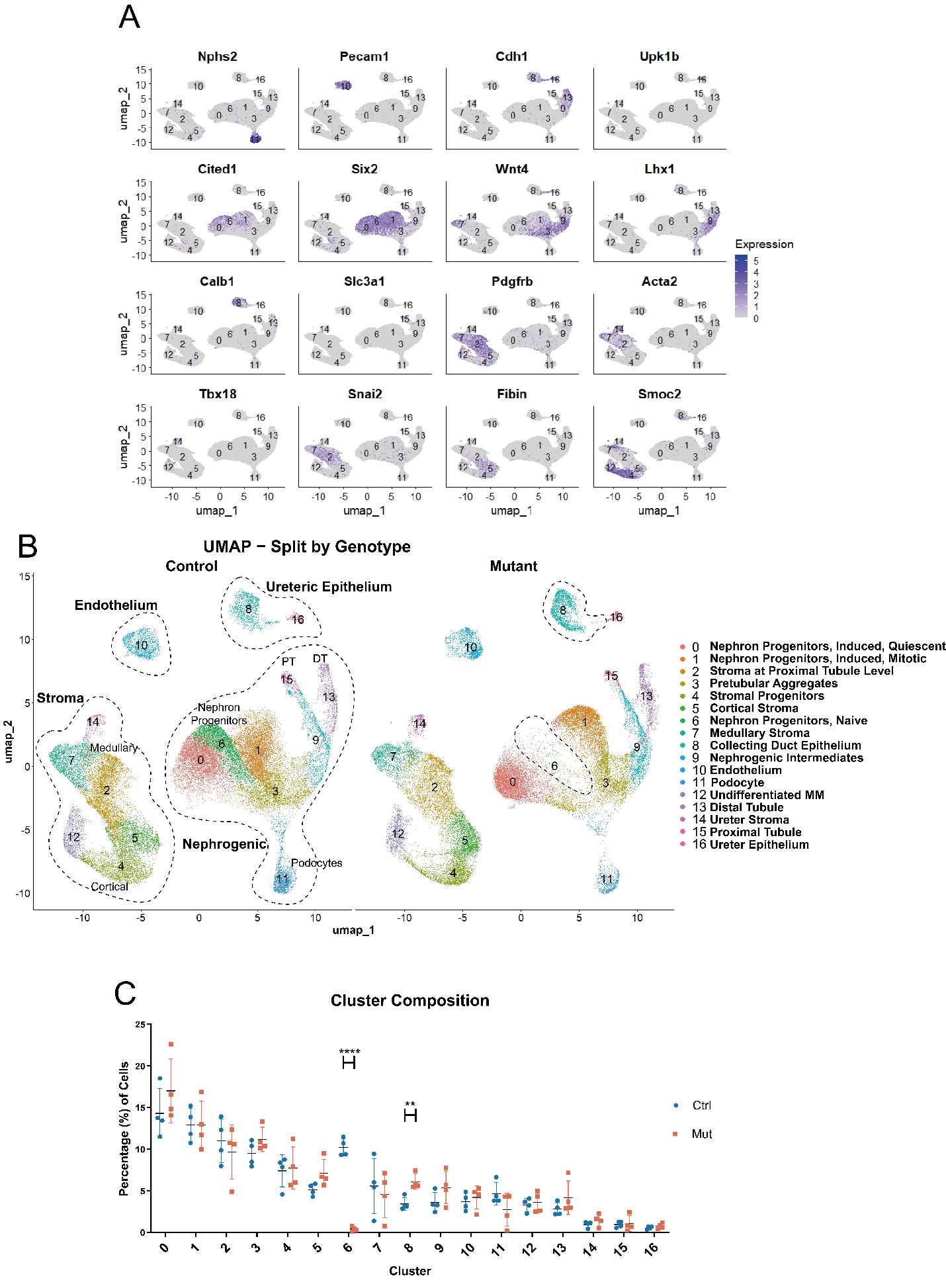

Supplemental Figure S3

(A) Expression of hallmark genes of kidney cell populations. (B) UMAP reduction of all cells that passed quality control screening in single-cell RNA sequencing, split by genotypes. Dashed lines in mutant panel highlights cell clusters with increased or decreased cells. (C) Cluster composition of each sample. Data are mean $\boldsymbol{\pm}$ SD. ** *p* < 0.01; **** *p* < 0.0001; (empirical Bayes moderated *t*-test).

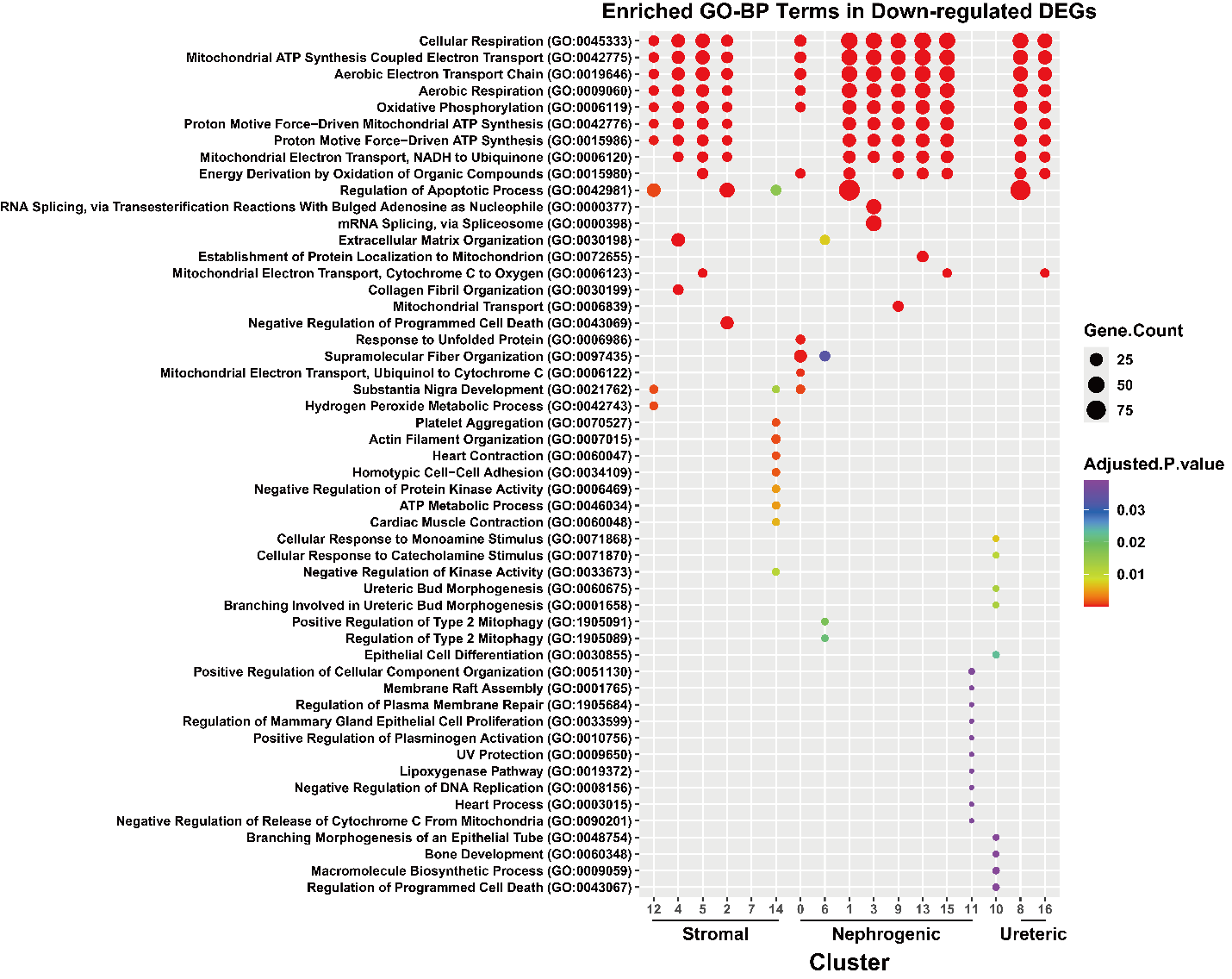

Supplemental Figure S4

Unbiased over-representation analysis on all pseudobulked kidney cell clusters. Terms were trimmed to show top 10 terms in each cluster ranked by adjusted *p* value.

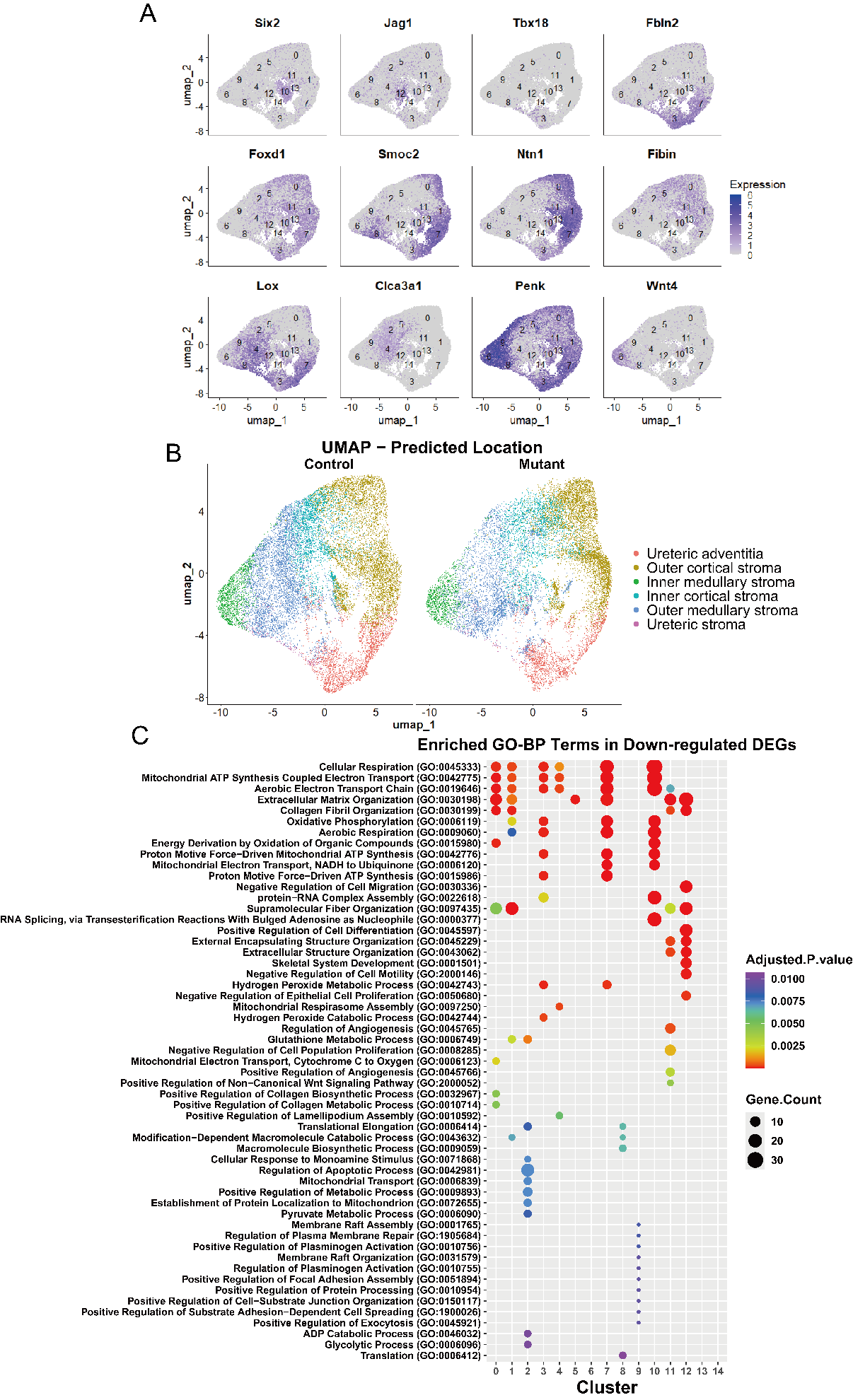

Supplemental Figure S5

Analysis of stromal subset of single-cell data. (A) Expression of hallmark genes that were used to define stromal heterogenity in E16.5 single-cell RNA sequencing. (B) Prediction of spatial location of each stromal cell by anchor transferring to a public spatial dataset (Jiang et al., 2025). “Inner medulla” labeled here corresponds to what was labelled as middle medulla in the original publication, since the original spatial dataset contains cells from E18.5 when a population closer to the papilla has emerged. (C) Unbiased over-representation analysis on all pseudobulked stromal cell clusters. Terms were trimmed to show top 10 terms in each cluster ranked by adjusted *p* value.

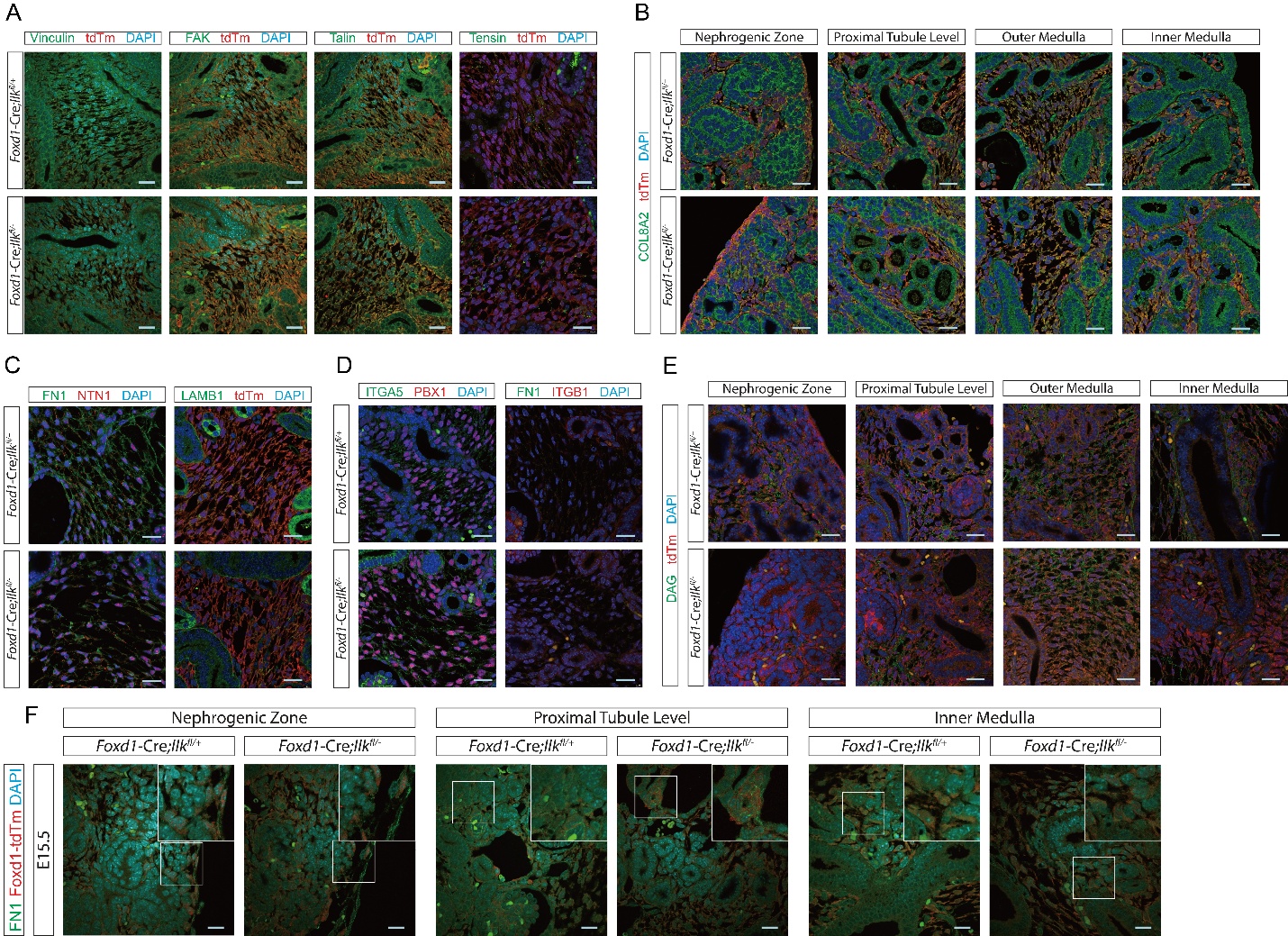

Supplemental Figure S6

(A) Immunostaining of focal adhesion components at E16.5. All tested components demonstrated comparable levels in control and mutant kidneys except vinculin. (B) Immunostaining of COL8A2 at E16.5. Images were captured across the corticomedullary axis. Mutant kidneys only demonstrated reduced expression in cortical stroma. (C) Immunostaining of fibronectin, netrin, and laminin at E16.5. Mutant kidneys only demonstrated disruption in fibronectin. (D) Immunostaining of ITGA5 and ITGB1 at E15.5. Expression was comparable in control and mutant kidneys. (E) Immunostaining of dystroglycan at E15.5. Images were captured across the corticomedullary axis. Mutant kidneys demonstrated decreased expression. However, the alteration was not detected in outer medulla. (F) Immunostaining of fibronectin at E15.5. No alteration was detected in the cortical zone, at the proximal tubule level, and in inner medulla. All scale bars represent 25 μm.

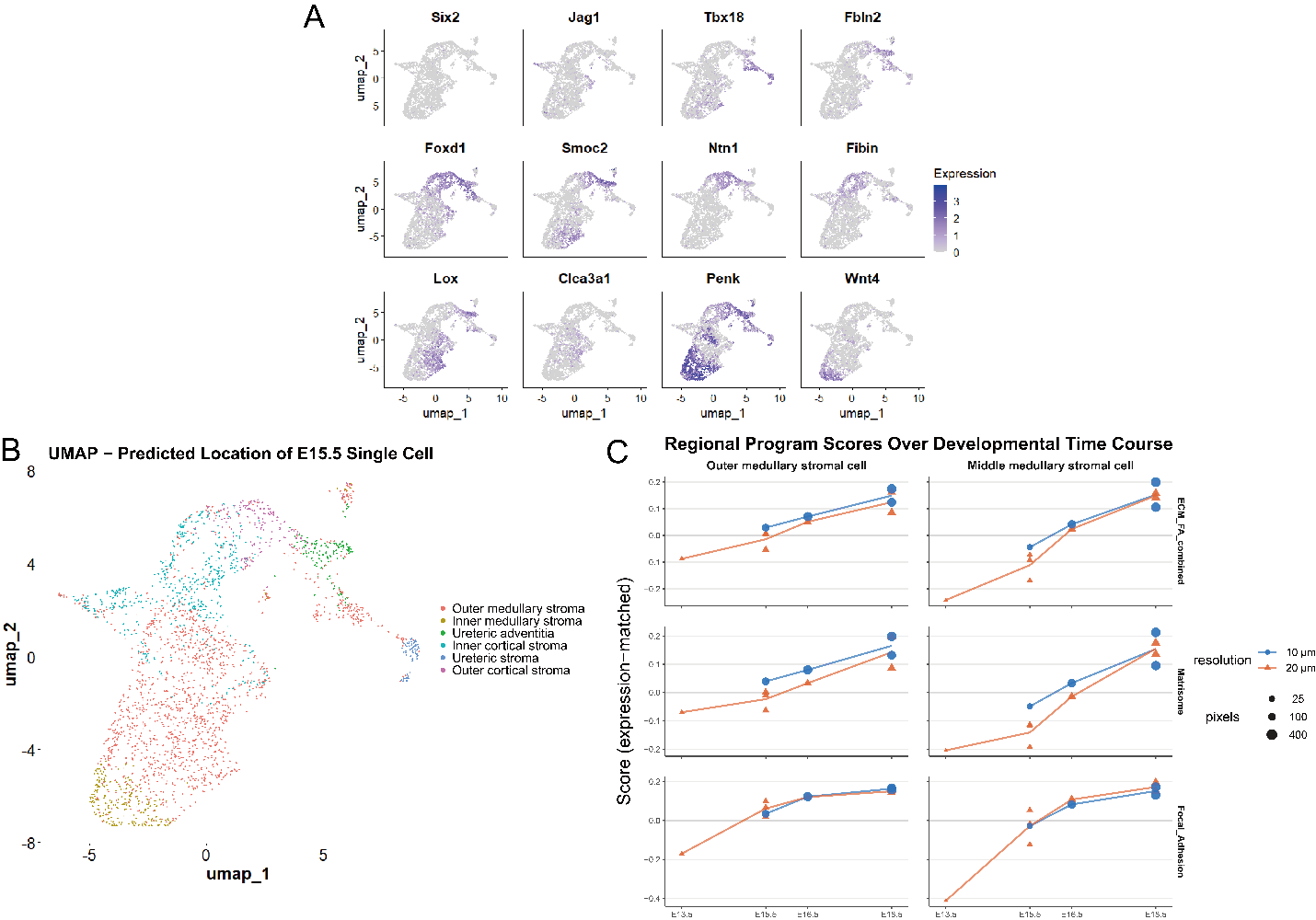

Supplemental Figure S7

(A) Expression of hallmark genes that were used to define stromal heterogenity in E15.5 public single-cell RNA sequencing datasets (Lawlor et al., 2019; Naganuma et al., 2021). (B) Prediction of spatial location of each E15.5 cell by anchor transferring to a public spatial dataset (Jiang et al., 2025). (C) Stroma pixels in time course spatial sequencing were scored for ECM and FA programs, both demonstrating an increase between E13.5 and E18.5. In two medullary regions, a change in slope was observed between E15.5 and E16.5 in both programs.
